# Dopaminergic Control of Retinal Oscillations Driving Infantile Nystagmus

**DOI:** 10.64898/2025.12.19.695412

**Authors:** B. Semihcan Sermet, Wouter Kamermans, Beerend H.J. Winkelman, Chris I. De Zeeuw, Maarten Kamermans

## Abstract

Infantile nystagmus is a debilitating involuntary eye movement disorder often associated with retinal diseases such as Congenital Stationary Night Blindness (CSNB). The oscillating eye movements of infantile nystagmus come with reduced visual acuity, strongly impairing quality of life. No cure exists for this condition. Previously, we demonstrated that nystagmus in the CSNB mouse model *Nyx^nob^* has a retinal cause. Specifically, we found that synchronized oscillations of retinal ganglion cells (RGCs) are transmitted to the accessory optic system, triggering compensatory eye movements. The RGC oscillations appear to originate from a specific retinal cell type, the A_II_ amacrine cell (A_II_ AC), making these cells the preferred target for treatment. Here we found that pharmacologically activating the dopaminergic input to A_II_ ACs in *Nyx^nob^* mice completely suppresses the pathological oscillations of both A_II_ ACs and RGCs and improves the signal fidelity of RGCs. Moreover, our retinal network simulations confirm that the experimentally observed changes to A_II_ AC voltage-gated currents are sufficient to account for the dopamine-dependent abolishment of these oscillations. Our findings provide a novel mechanistic understanding of the retinal mechanism underlying infantile nystagmus as well as the associated low visual performance. Consequently, they offer the first pharmacological therapeutic strategy for this disorder.

**Significance:** Oscillatory eye movements in infantile nystagmus arise from abnormal retinal activity, yet the cellular basis of this instability has remained unclear. Here we show that this abnormal activity stems from disrupted dopaminergic modulation. By defining how dopamine regulates the activity of a key retinal cell type, the A_II_ amacrine cell, we demonstrate that restoring dopamine levels can return the retinal circuit to a stable state and improve the clarity of visual signals leaving the eye. This work links neuromodulation to retinal circuit instability and identifies dopaminergic control of A_II_ amacrine cells as a pharmacologically targetable point of intervention. These findings suggest a retina-focused pharmacological treatment strategy for infantile nystagmus, a disorder that currently lacks effective therapeutic options.

## Introduction

One of the remarkable abilities of our brain is to stabilize images on the retina while moving around in our environment and, by doing so, facilitate perception. This task is accomplished by integrating signals from the retina and vestibular hair cells within the neural circuits controlling eye movements, including the accessory optic system (AOS) and the vestibulocerebellum (1). These systems, in turn, modulate the activity of abducens motor neurons, which drive the eye muscles. Multiple feedback loops and adaptive mechanisms act to minimize image movement over the retinal surface, i.e., retinal slip. When these mechanisms malfunction or are misaligned, nystagmus can be a consequence (2).

There are many forms of nystagmus (prevalence: approx. 0.2%) with many different causes. Nystagmus can be infantile or develop later in life, but in either case it is often accompanied by decreased visual acuity (3–6). This also holds for patients suffering from congenital stationary night blindness (CSNB), which is caused by genetic mutations affecting transmission at the photoreceptor to ON-bipolar cell (ON-BC) synapse (7, 8). Some CSNB patients with infantile nystagmus have a mutation in the retina-specific protein nyctalopin (9). By analyzing the so-called *Nyx^nob^* mouse model, which expresses non-functional nyctalopin (10), we have been able to show that malfunction of the photoreceptor to ON-BC synapse makes retinal ganglion cells (RGCs) oscillate (11–13). These oscillatory signals are sent to the brain areas responsible for image stabilization, which in turn results in oscillatory eye movements, i.e., infantile nystagmus.

Previous work strongly implicates A_II_ amacrine cells (A_II_ ACs) as the source of this oscillatory firing of RGCs (12, 13). A_II_ ACs are highly connected neurons in the inner retina (Figure S1); they are electrically coupled with each other as well as with cone ON-BCs, and they provide glycinergic inhibition to cone OFF-BCs, other ACs, and RGCs (14). In the dark-adapted retina, A_II_ ACs receive rod signals from the rod ON-BCs, amplify these signals by an active mechanism involving voltage-gated ion channels (I_Na_, I_K-A_, I_K-M_) (15) and transmit them to the ON- and OFF-pathways (16). In the light-adapted retina, the amplification by A_II_ ACs is not needed because now the large cone signals are transmitted directly from the BCs to the RGC, suggesting that the properties of A_II_ ACs are modulated during light/dark adaptation, most likely by dopamine.

This active amplification in the A_II_ ACs depends on the specific cellular arrangement of the ion channels. I_Na_ and I_K-M_ are exclusively localized on a highly specialized process of the A_II_ AC, the initial segment (17) (Figure. 2A), while I_K-A_ is present in the soma as well (18). Under pathological circumstances, this amplification process can become unstable and lead to intrinsic oscillations, turning the A_II_ AC into an intrinsic oscillator (19, 20, 11).

Dopamine (DA) is well-positioned to be the modulator of A_II_ AC amplification. DA levels change during the circadian cycle, with DA levels high in the light-adapted retina and low in the dark-adapted retina (21). A_II_ ACs receive a D1 dopaminergic input from dopaminergic amacrine cells (DAC) (22–25, 17), suggesting the activity of their voltage-gated ion channels may be modulated by DA. Indeed, Veruki and colleagues (2025) showed evidence of the dopaminergic downregulation of the sodium channels of A_II_ ACs in rats, leading to a shift in spike threshold. This implies that in the dark-adapted retina, low levels of DA allow for highly activated A_II_ AC voltage-gated ion channels leading to maximal amplification, enabling high-fidelity transmission of the rod signals. Conversely, in light-adapted conditions, there is no need for amplification of the large cone signals, and the increased DA levels may reduce the sensitivity of the ion channels at the initial segment, minimizing the amplification.

In CSNB, DA release is reduced (26, 27), consistent with the notion that the activity of DACs is modulated by the rod-driven ON pathway (28, 29). Thus, we hypothesize that because of the low DA concentration in CSNB, the voltage-gated ion channels on the initial segment of the A_II_ AC remain highly active in the light-adapted state and lead to spontaneous oscillations propagating in the retinal network, ultimately driving oscillating activity of RGCs and the oculomotor system downstream (12, 13).

## Results

### DA blocks RGC and A_II_ AC oscillations

To test the hypothesis that DA modulates the oscillations seen in RGCs, we recorded spontaneous RGC activity from isolated light-adapted retinas of *Nyx^nob^* and wild-type (WT) mice using a multielectrode array (MEA). As shown by spike raster plots, Fourier transform, and autocorrelation analysis, many RGCs of *Nyx^nob^* mice, but not of WTs, exhibited robust spontaneous oscillations with a dominant frequency around 5 Hz (Figure 1B, E, and F). Application of 10 µM DA to the *Nyx^nob^* retina suppressed these oscillations (Figure 1C). Quantitative analysis of mean power spectral density confirmed strong oscillations in baseline RGCs (mean 1.44 × 10^−3^ sps^2^/Hz, n = 165), which were significantly reduced by DA application (Figure 1G; mean 0.124 × 10^−3^ sps^2^/Hz, n = 165; *P* < 0.0001, Wilcoxon matched-pairs signed rank test), bringing the power of the dominant frequency to a level comparable to that of WT. Moreover, the DA effect was reversible in that a washout for 30 mins allowed the oscillations to reoccur (Figure 1F). To identify the type of DA receptor mediating this suppression, we tested the specific D1 DA agonist SKF-81297. Application of 10 µM SKF-81297 was also highly effective in suppressing these oscillations (Figure 1D, F, and H), reducing mean power to 0.06 × 10^−3^ sps^2^/Hz (n = 88 RGCs; *P* < 0.0001 vs baseline, Wilcoxon matched-pairs signed rank test). The efficacy of SKF-81297 was very high, indicated by a statistically significant difference when compared directly to DA (*P* = 0.0126, DA vs SKF, Mann-Whitney test). In contrast, application of the D2-agonist quinpirole hydrochloride (QPH, 10 µM) failed to eliminate RGC oscillations (Figure S2B, C), indicating that D1 receptor activation is the primary mechanism for suppressing this pathological activity.

**Figure 1.**
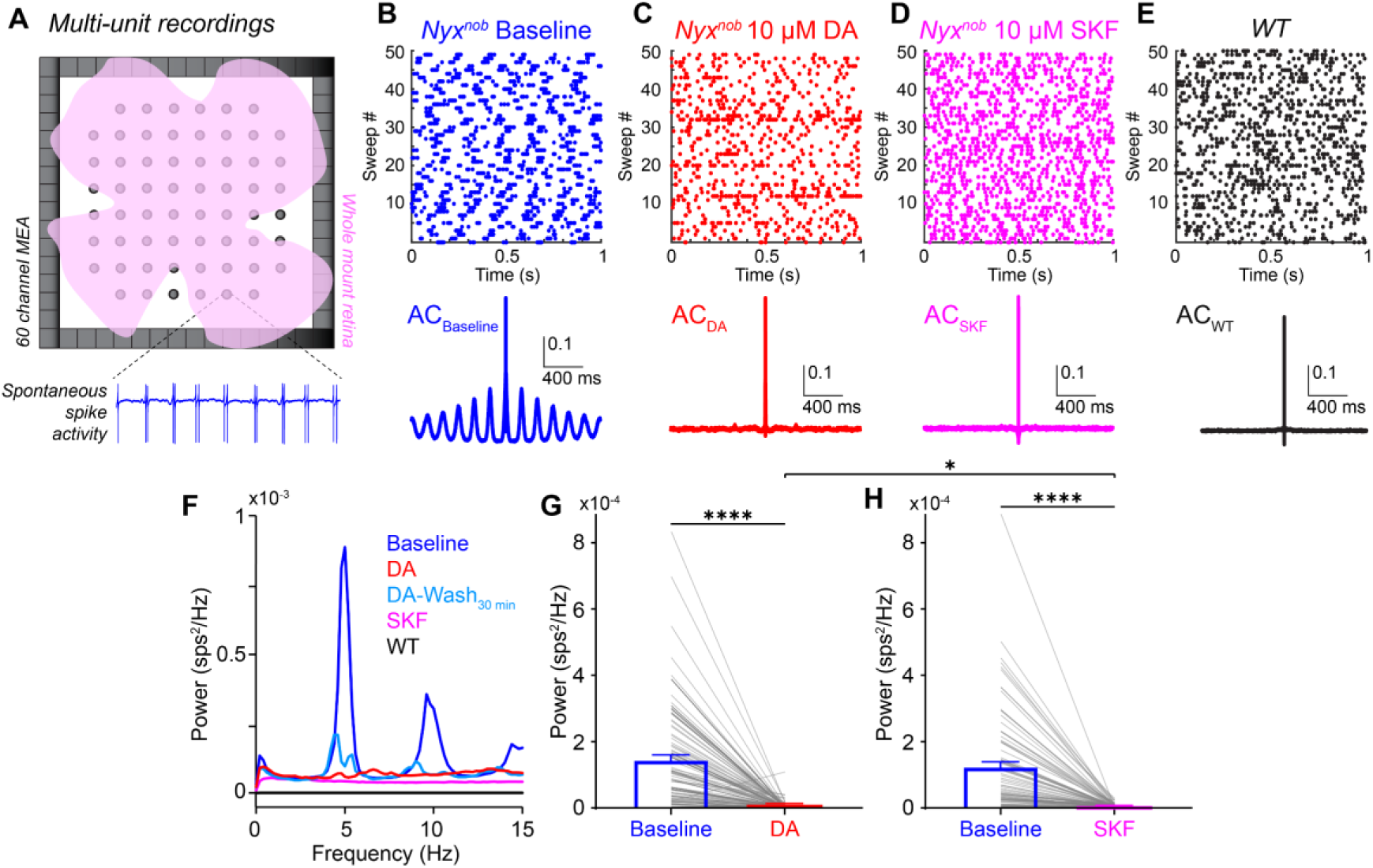
Dopamine suppresses retinal ganglion cell oscillations ex vivo in *Nyx^nob^* mice. **A**, Schematic of whole-mount retina recording on a multielectrode array (MEA). **B-E**, Spontaneous retinal ganglion cell (RGC) activity in the dark in an isolated light-adapted retina. Top: raster plot of a single RGC. Bottom: autocorrelation of the same RGC. **B**, Baseline *Nyx^nob^* mouse retina showing robust oscillations. **C**, *Nyx^nob^* retina with 10 µM dopamine (DA) application. **D**, *Nyx^nob^* retina with 10 µM D1 agonist SKF-81297 (SKF) application, showing complete suppression of oscillations. **E**, Wild-type (WT) mouse retina showing no spontaneous oscillations. **F**, Mean power spectral density of RGC activity from a representative retina under baseline conditions (blue), with DA (red), after DA washout (cyan), with SKF (magenta), and in a WT retina (black). **G**, Paired comparison of baseline vs DA oscillation power for individual RGCs (gray lines), with mean ± SEM overlaid (y-axis in sps^2^/Hz, sps = spikes per second). Baseline (blue, mean = 1.44 × 10^−3^ sps^2^/Hz, n = 165), DA (red, mean = 0.12 sps^2^/Hz × 10^−3^, n = 165). Baseline vs DA, *P* < 0.0001 (Wilcoxon paired test). **H**, Same analysis as in G, but for baseline (blue, mean = 1.23 × 10^−3^ sps^2^/Hz, n = 88) vs SKF (magenta, mean = 0.06 × 10^−3^ sps^2^/Hz, n = 88). Baseline vs SKF, *P* < 0.0001 (Wilcoxon paired test). DA vs SKF, *P* = 0.013 (Mann-Whitney test).

Since the RGC oscillations appear to originate from A_II_ ACs (12, 13), we next performed whole-cell patch-clamp recordings from A_II_ ACs (Figure 2A) in light-adapted retinal slices from *Nyx^nob^* mice. Application of 10 µM DA did not significantly change their resting membrane potential (Figure S3). To understand the ion-channel basis, we examined voltage-gated currents using voltage-clamp recordings from A_II_ ACs. Voltage steps elicited large transient outward, transient inward, and sustained currents in baseline *Nyx^nob^* A_II_ ACs (Figure 2B). The transient outward and inward currents most likely consist of the A-type potassium (I_K-A_) and a sodium current (I_Na_), respectively (20, 30). Instead, the sustained currents may reflect a combination of the I_K-A_ and M-type potassium (I_K-M_) currents. At a clamp potential of −50 mV, many unclamped spike-like events (spikelets) were observed (Figure 2B). The fact that we were unable to clamp these spikelets suggests that they are initiated at the initial segment, a compartment connected to the soma via a high-resistance cable-like structure (17). This means that the recordings suffer from a space clamp problem and that the IV relations for especially I_K-M_ and I_Na_ are rough estimates of the IV relations of these currents. Application of 10 µM DA visibly reduced the amplitude of I_K-A_ and I_Na_ currents of the A_II_ ACs in *Nyx^nob^* mice, while the amplitude of the sustained current (I_K-M_) did not change significantly in the physiological membrane potential range (Figure 2B). Current-voltage (IV) relationships confirmed a significant reduction in the peak amplitude of both I_K-A_ and I_Na_ currents by DA (Figure 2C, D). Moreover, we observed a noticeable reduction in the number of spikelets (Figure 2B).

**Figure 2.**
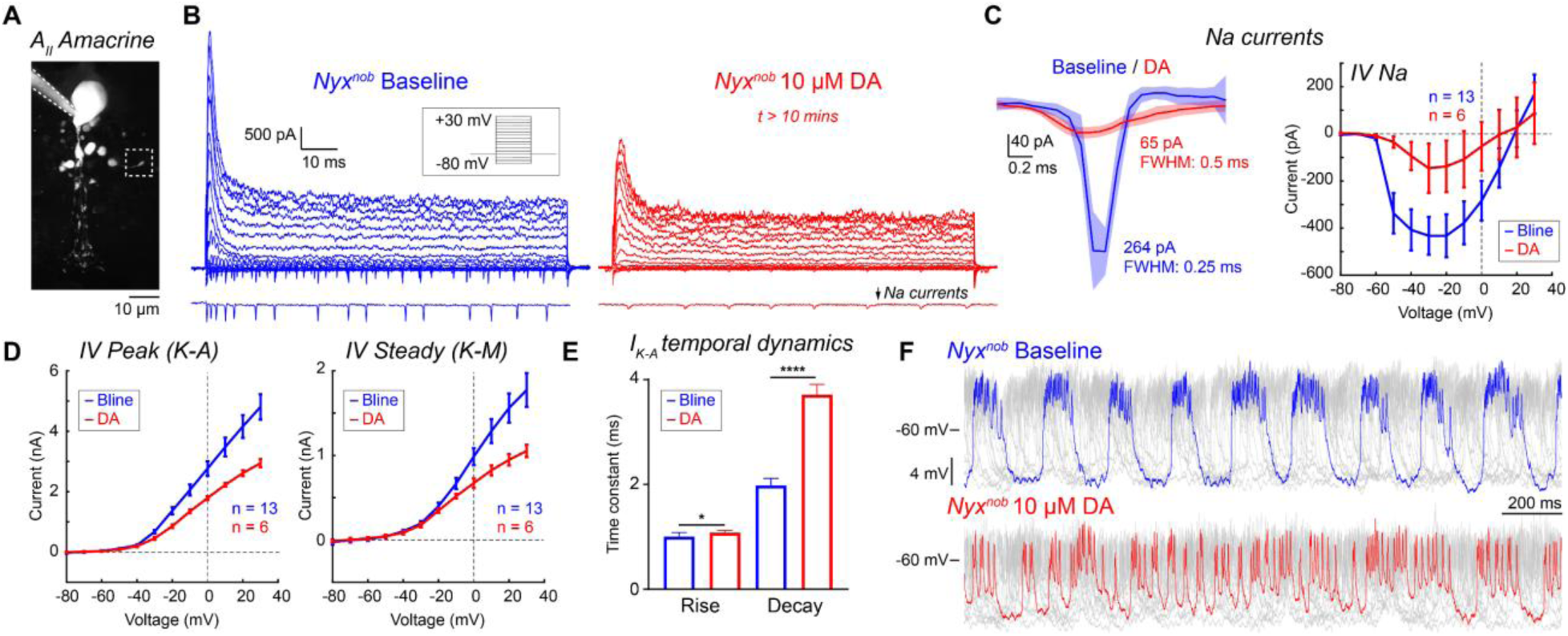
Dopamine modulates voltage-gated currents and suppresses oscillations in *Nyx^nob^* A_II_ amacrine cells. **A**, Two-photon z-stack of a Lucifer Yellow-filled A_II_ AC in a retinal slice during whole-cell recording. The initial segment is highlighted with a white dashed square **B**, Left: Voltage-clamp currents from a *Nyx^nob^* A_II_ AC (from −80 mV to +40 mV in 10 mV increments from a holding potential of −60 mV). Bottom: Isolated −50 mV step response, highlighting transient inward currents (I_Na_) and unclamped spikelets. Right: After 10 µM dopamine (DA) application, showing reduced current amplitudes and spikelet activity. **C**, Dopaminergic modulation of I_Na_ and spikelet kinetics. Left: Isolated mean I_Na_ traces from a representative A_II_ AC in baseline (blue) and with DA (red). Right: Mean I-V for peak I_Na_ in baseline (blue) and DA (red) conditions. **D**, Mean I-V relationships for estimated potassium currents. Left: Peak outward current (0-10 ms after stimulus onset, mainly I_K−A_) in baseline (blue) and DA (red) conditions, showing DA-induced reduction. Right: Sustained outward current (80-100 ms after stimulus onset, mainly I_K-M_), in baseline (blue) and DA (red) conditions, no significant DA effect in physiological range. **E**, Bar graphs of I_K−A_ temporal dynamics. Rise times: baseline (blue; mean 1.00 ms, n = 13) vs DA (red; mean 1.08 ms, n = 6; *P* = 0.016, Mann-Whitney test). Decay time constants: baseline (blue; mean 1.98 ms, n = 13) versus DA (red; mean 3.71 ms, n = 6; *P* < 0.0001, Mann-Whitney test). DA significantly alters both rise and decay kinetics of I_K-A_. **F**, Current-clamp recordings from a *Nyx^nob^* A_II_ AC. Top: Baseline (blue): ~5 Hz membrane potential oscillations with steady hyperpolarizing current.. Bottom: Application of 10 µM DA (red) disrupts oscillations, leading to an irregular membrane potential. All summary data are presented as mean ± SEM.

When we further analyzed the kinetics of the voltage-gated currents (Figure 2E), DA application turned out to induce a small, but statistically significant, increase in the rise time for I_K−A_ in the A_II_ ACs of *Nyx^nob^* mice (baseline: 1.00 ms, n = 13 cells vs DA: 1.08 ms, n = 6 cells; *P* = 0.0156, Mann-Whitney test). More prominently, DA significantly slowed the decay kinetics of I_K−A_, nearly doubling the decay time constant (baseline: 1.98 ms, n = 13 cells vs DA: 3.71 ms, n = 6 cells; *P* < 0.0001, Mann-Whitney test). In addition, DA seemed to have an even more pronounced effect on I_Na_ kinetics (Figure 2C). It reduced the estimate of the peak I_Na_ amplitude fourfold (Figure 2C). Moreover, the rise time and decay time of the spikelets increased about two-fold and five-fold, respectively (Figure 2C). These findings indicate that DA profoundly modulates A_II_ AC excitability not only by reducing the magnitude of key voltage-gated currents (I_Na_ and I_K-A_) but also by significantly slowing their activation and inactivation kinetics, rendering the A_II_ ACs less prone to oscillations and reducing the number of spikelets.

The data highlighted above indicate that the combination of voltage-gated channels in the initial segment produces the spikelets and that this active compartment, in combination with the sustained voltage-gated potassium current (I_K-M_), makes the A_II_ AC able to generate large oscillations with spikelets that are superimposed on these oscillations. Moreover, this configuration implies that the A_II_ ACs operate, depending on their membrane potential, in a spiking regime, an oscillatory regime, or a graded potential regime. To further establish the impact of DA on the oscillations of the A_II_ ACs, we further investigated their relation with the membrane potential. Under baseline conditions with a steady hyperpolarizing current injection, *Nyx^nob^* A_II_ ACs display membrane potential oscillations around 5 Hz (Figure 2F). At the most depolarized phase of the oscillations, spikelets were visible. Application of 10 µM DA disrupted these oscillations, resulting in an irregular membrane potential and irregular spikelets (Figure 2F), showing that DA reduces the cell’s propensity to oscillate. Together, these experiments show that DA modulates the ion channels that constitute the intrinsic oscillator in the A_II_ ACs. At low DA concentrations, A_II_ ACs in *Nyx^nob^* can oscillate, whereas at high concentrations, the oscillations are suppressed.

### A_II_ AC ion channel modulation is sufficient to block oscillations

Since DA might also have modulated other retinal cell types (21), we wanted to assess in a computational model (11) to what extent the impact of DA on the properties of the voltage-gated currents of the A_II_ ACs is sufficient to explain the observed effects on the oscillatory behavior. The model we developed for *Nyx^nob^* A_II_ ACs reproduced both the voltage step responses and the IVs of the transient and sustained currents under baseline conditions (Figure 3A-D). Moreover, reducing the conductance, increasing the inactivation time constant, and reducing the activation/inactivation ratio of the I_K-A_ was sufficient to simulate the DA condition (Figure 3B; Table 1). However, it should be noted that these latter changes did not affect the spikelets; the spikelets are generated by the combined activity of I_Na_, I_K-A_, and I_K-M_. Since I_Na_ and I_K-M_ are located in the initial segment, we don’t have reliable direct measurements of these currents, and we had to estimate the related parameters (Figure 3D; see also Figure 2C, D). To reproduce the spikelet response in the DA condition, it was necessary to reduce the conductance and increase both the activation and inactivation time constants of I_Na_ (Table 1).

**Figure 3.**
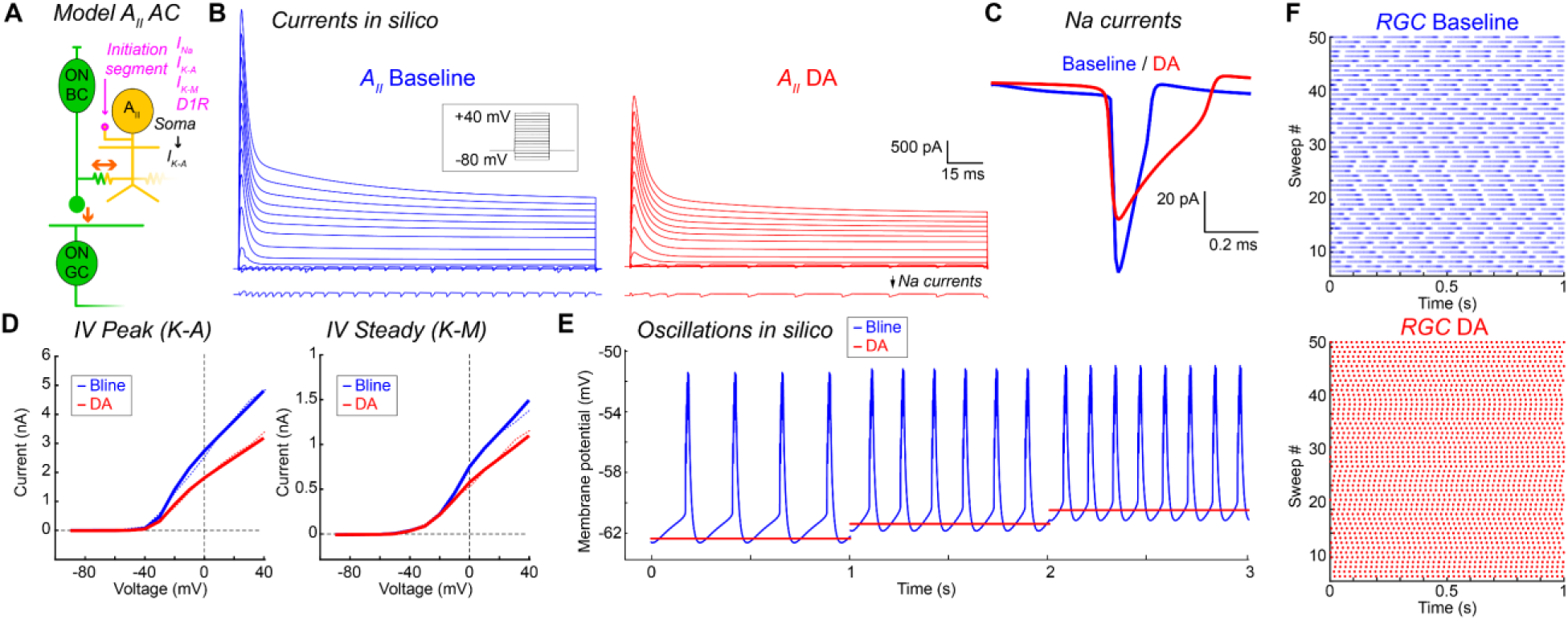
Computational model of a retinal microcircuit recapitulates dopaminergic modulation of excitability and oscillations. **A,** Schematic of the model consisted of an ON bipolar cell (ON BC), an A_II_ amacrine cell (A_II_ AC), and an ON ganglion cell (ON GC), showing voltage-gated sodium current (I_Na_), A-type potassium current (I_K−A_), M-type potassium current (I_K−M_), and the D1 receptor (D1R) at the initial segment, and a somatic I_K−A_. **B,** Simulated A_II_ AC voltage-clamp responses to depolarizing voltage steps (−80 mV to +40 mV). Left: Baseline (blue traces). Right: Simulated effect of dopamine (DA model) application (red). Bottom traces show isolated responses to a −50 mV step, highlighting simulated I_Na_-mediated spikelets and their reduction by DA. **C,** Isolated simulated transient inward sodium currents (I_Na_) for baseline (blue) and DA (red) conditions, showing a DA-induced amplitude reduction. **D,** Model I-V relationships. Left: Peak inward (I_Na_) and outward (I_K−A_) currents. Right: Steady-state outward current (I_K−M_). Solid lines are model simulations, and dashed lines are experimental data from a single A_II_ AC for comparison. **E,** Simulated current-clamp behavior of the A_II_ AC model over 3 seconds with incrementally increasing current injection. Baseline (blue) shows oscillations at approximately 5-9 Hz with increasing stimulus. In the DA condition (red), the model remains stable at lower current injections, until it oscillates at a relatively low frequency (~3 Hz) at the highest current injection. Modeled dopaminergic modulation, thus significantly stabilizes the A_II_ AC, protecting against spontaneous oscillations. **F,** Top: raster plot of the model RGC in baseline condition oscillating at approximately 7 Hz. Bottom: same RGC in DA condition without oscillatory activity.

**Table 1.**
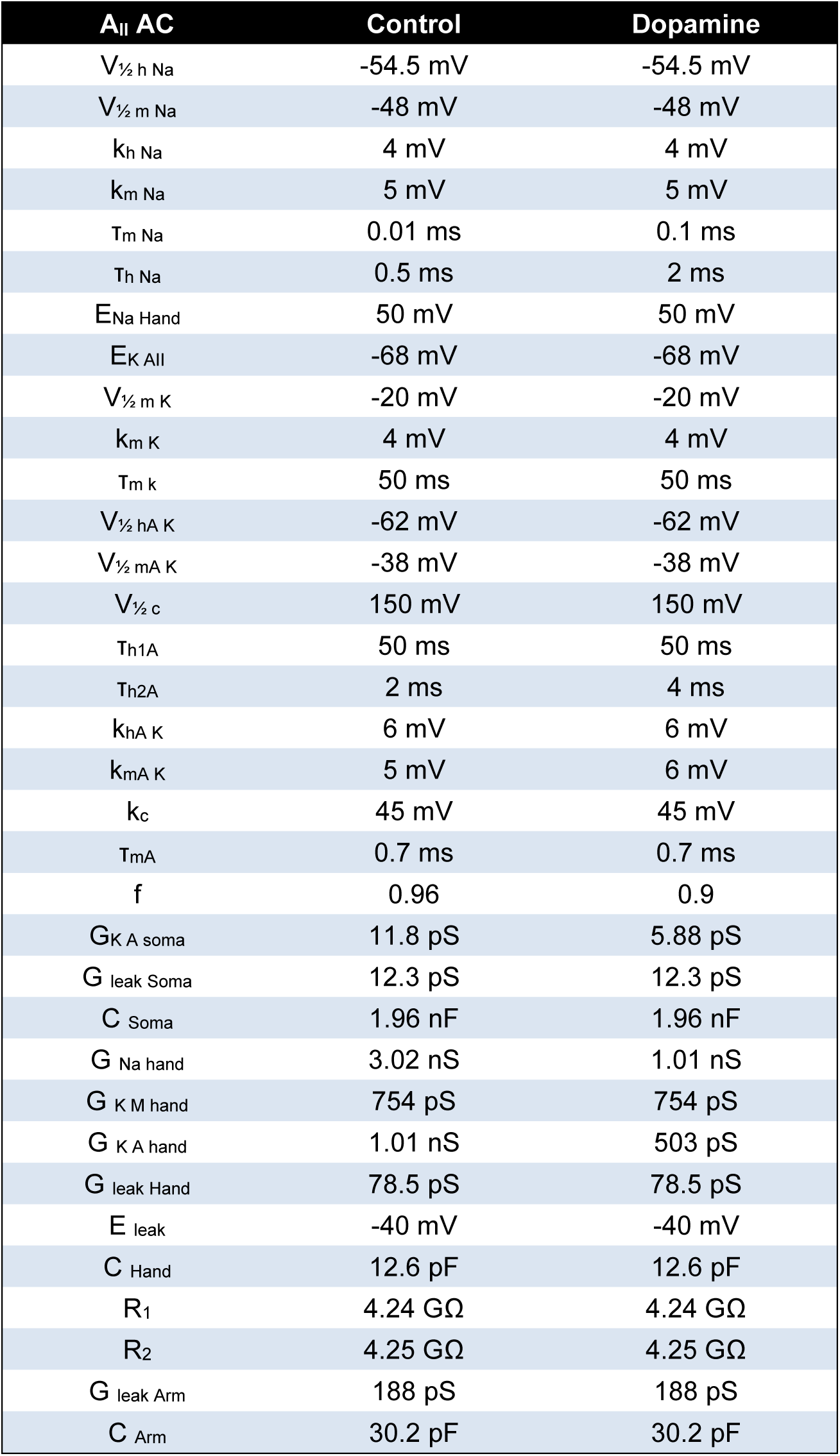
A_II_ AC model parameters.

Next, we tested how the oscillatory behavior of the A_II_ ACs is affected by these changes in model parameters. The computational model shows that the modulation of the ionic currents, as described above, has a pronounced effect on the oscillatory behavior of the A_II_ ACs (Figure 3E). Whereas the baseline model exhibited robust oscillations that increased in frequency (e.g., from ~5 Hz to ~9 Hz) with increasing current injection, the DA-modulated model remained stable over almost the whole current injection range. Only with the strongest current injection, some bursting activity occurred. Critically, the model also predicts the effect on retinal output. As shown by the modeled RGC activity (Figure 3F), the baseline condition shows oscillatory activity approximately at 7Hz. However, with A_II_ AC specific DA modulation, the model predicts the complete abolishment of the oscillations (Figure 3F, bottom). These simulations show that the changes we have determined experimentally in the properties of the voltage-gated currents of A_II_ ACs are sufficient to account for the DA-dependent abolishment of their pathological oscillatory behavior in the *Nyx^nob^* mice.

### DA enhances RGC light response fidelity

The spontaneous oscillations in *Nyx^nob^* RGCs could potentially interfere with the encoding of visual information by increasing the “noise”. To investigate this, we compared RGC responses to a 500 ms light flash in baseline conditions and in the presence of 10 µM DA or SKF-81297 (Figure 4). In baseline *Nyx^nob^* retinas, the RGC activity in the dark was strongly oscillatory, whereas the light responses of the RGCs were often very small (Figure 4B, D). However, upon application of DA, the background oscillations were suppressed, and in addition, the light responses of the RGCs were significantly larger (Fig. 4C, E). In this example, the peak light response amplitude increased approximately fourfold after application of DA (Figure 4D, E). To further quantify this effect, we first measured the standard deviation (SD) of the baseline firing rate of the RGC activity during the dark period preceding the light flash. All individual traces were aligned such that the oscillations were in phase. We consider the SD measured this way as noise. Figures 4F and 4H show that the SD before and after DA or SKF-81297 applications was significantly reduced (*P* < 0.0001 for both, *SD_Bline_* = 7.3, *SD_DA_* = 5.7, *SD_SKF_* = 5.1, Wilcoxon test), with no significant difference between DA and SKF-81297. Instead, application of the D2-agonist QPH had no effect on the SD of the baseline firing (Figure S2F).

**Figure 4.**
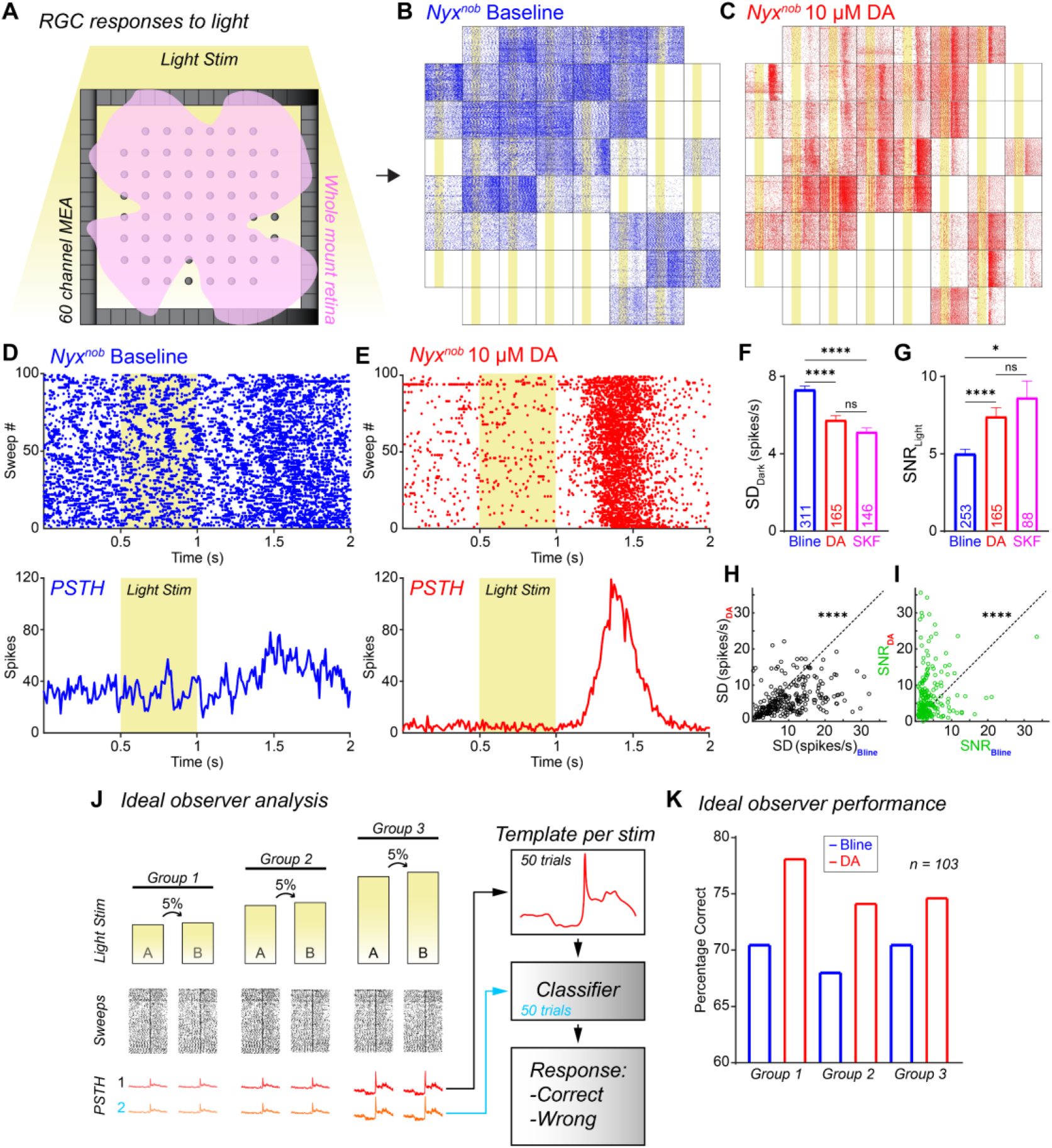
Dopamine enhances retinal ganglion cell light response fidelity in *Nyx^nob^* mice. **A**, Schematic of retina MEA recording with a 500 ms light stimulation, repeated every 2 seconds. **B**,**C**, Global RGC activity raster plots. **B**: Baseline *Nyx^nob^* retina (blue), with oscillatory background activity and modest light responses. **C**: Same *Nyx^nob^* retina after 10 µM dopamine (DA, red), showing suppressed background oscillations and enhanced light responses. **D**, Single RGC’s response to light stimulation (baseline). Top: Raster plot (blue). Bottom: Corresponding peri-stimulus time histogram (PSTH). **E**, Same RGC as in (**D**) after 10 µM DA (red), showing reduced baseline activity and increased peak light response. **F**, Mean standard deviation (SD) of baseline RGC firing rates (noise) under baseline (blue), 10 µM DA (red), and 10 µM SKF-81297 (SKF; magenta). Both DA and SKF significantly reduce baseline noise (*P* < 0.0001, Wilcoxon test; see text for values). **G**, Mean signal-to-noise ratio (SNR) for baseline (blue), 10 µM DA (red), and 10 µM SKF (magenta). Both DA and SKF significantly increase SNR (*P* < 0.0001 for DA, *P* = 0.018 for SKF, Wilcoxon test; see text for values). **H**, **I**, Scatter plots of paired comparisons for individual RGCs. h: Baseline SD after DA versus baseline (SD_DA_ vs SD_Bline_; black circles). i: SNR after DA versus baseline (SNR_DA_ vs SNR_Bline_; green circles). All summary data in bar graphs are mean ± SEM. **J**, Schematic of ideal observer analysis. Masks are generated from the PSTH of half the trials (50 trials). A classifier then matches the PTSH of the other half of the responses to the masks (Correct/Wrong). This measures discriminability across three intensity groups. **K**, Percentage Correct classification under baseline (Bline, blue) and 10 µM DA (DA, red). DA consistently increases correct classifications across all groups, demonstrating enhanced RGC discrimination (n = 103 RGCs).

Next, we determined the amplitude of the peak offset light-response for all RGCs without further classification. This is a crude measure for the amplitude of the light-response of the whole retina, further referred to as the signal. We calculated the signal-to-noise ratio (SNR) by dividing the signal by the SD of the dark activity for each RGC recorded from. Both DA and SKF-81297 significantly increased the SNR compared to the baseline condition (Figure 4G, I, *P* < 0.0001 for DA, *P* = 0.018 for SKF, *SNR_Bline_* = 5.0, *SNR_DA_* = 7.4, *SNR_SKF_* = 8.6, Wilcoxon test). Interestingly, application of the D2-agonist QPH also led to a significant increase in the SNR (Figure S2G, P < 0.0001 for QPH, *SNR_Bline_* = 5.4, *SNR_QPH_* = 9.4, Wilcoxon test). Since the baseline noise was not significantly affected by QPH, this improvement in SNR must be attributed to an increase in the signal.

Finally, we determined how dopamine affected the discriminative power of RGCs by using an ideal observer approach (Figure 4J). Hundred responses to two flash stimuli of almost equal intensity (5% difference) were recorded on the MEA before and after the application of dopamine. This dataset was used to train and test the ideal observer. Specifically, fifty responses (odd-numbered trials) were used to generate a template PSTH for each recorded unit, and the remaining fifty responses (even-numbered trials) were used as a dataset for the classifier. For each test response, the sum of squared errors between the templates for stim A and B and the dataset was calculated. The lowest sum of squared errors was used to classify the response to be most likely generated by stimulus A or B. In this way we could generate a histogram for the detection of either stimulus A or B. Figure 4K shows that application of dopamine markedly increases the discriminative power of RGCs. Thus, our data show that DA application in *Nyx^nob^* mice does not only suppress the retinal oscillations but also improves the signal quality of RGCs by lowering the noise via D1 receptors and increasing the signal via both D1 and D2 receptors.

## Discussion

In this paper we show that activation of D1 DA receptors in the retina of *Nyx^nob^* mice reduces the sensitivity of A_II_ ACs by affecting their I_KA_ and I_Na_ channels and thereby their propensity to oscillate (Figure 5). As a consequence, DA modulation effectively blocks oscillations of the A_II_ ACs, preventing oscillatory activity from propagating to the RGCs and from there to the AOS. As we have shown a causal relation between RGC oscillations and nystagmus (11–13), our data open up the avenue to exploit dopaminergic drugs as a treatment for infantile nystagmus.

**Figure 5.**
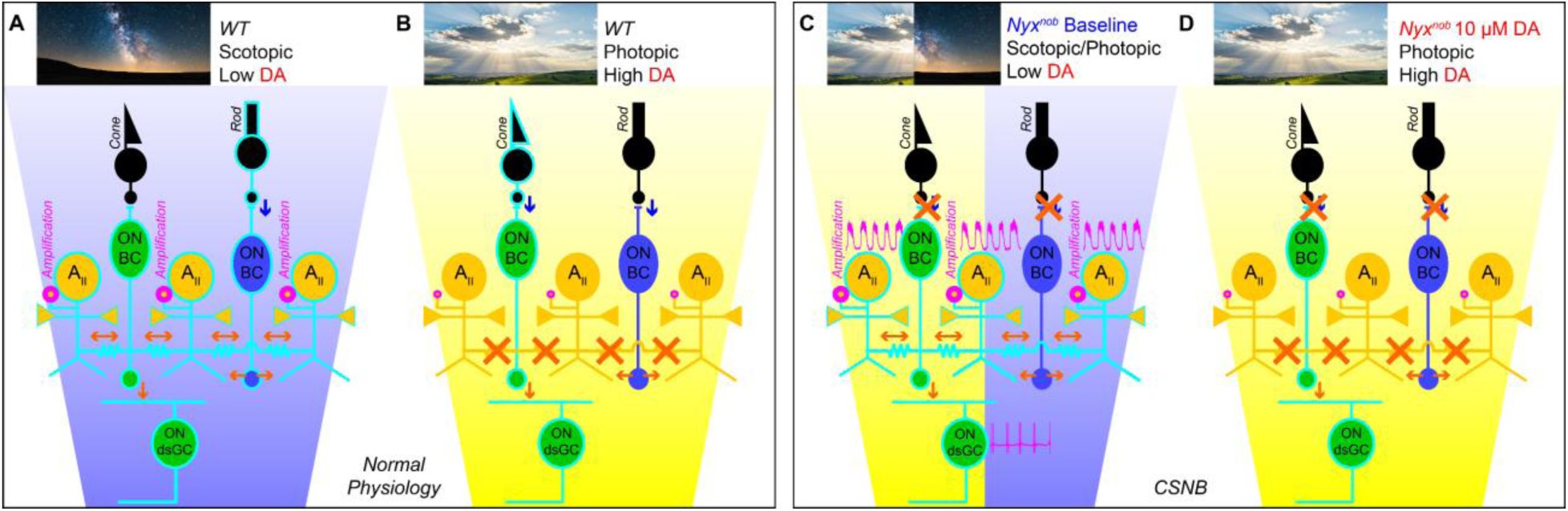
Schematic summary of dopaminergic control of A_II_ amacrine cell activity and its implications for nystagmus in CSNB. The A_II_ amacrine cell (AC) microcircuit depicts a cone and a rod providing input to their respective ON bipolar cells (ON-BCs). These ON-BCs connect to A_II_ ACs, interconnected by gap-junctions and provide input to ON retinal ganglion cell (ON-RGC) including the ON directional RGCs (ON-dsRGCs). Signal flow is indicated by arrows: blue are sign-inverting, and red are sign-conserving pathways. Active cells and pathways are highlighted. **A**, Scotopic (Low DA): rods transmit light signals, relayed via rod ON-BCs to A_II_ ACs. A_II_ ACs are electrically coupled to each other and cone ON-BCs, and their initial segments are highly sensitive, amplifying weak rod signals for transmission to RGCs. **B**, Photopic (High DA): Dopamine levels are high, cones transmit light signals, activating cone ON-BCs. High DA levels reduce the coupling between A_II_ ACs and desensitize their initial segments, preventing unnecessary amplification of robust cone signals and ensuring stable retinal output. **C**, Photopic or scotopic conditions in CSNB (*Nyx^nob^*): The genetic defect disrupts the photoreceptor to ON-BC synapse, leading to a dysfunctional ON-pathway and pathologically low retinal DA levels. Consequently, the retina remains in its scotopic wiring even in the photopic condition. A_II_ ACs remain coupled, and their initial segments remain hypersensitive. This renders A_II_ ACs unstable, causing them to generate spontaneous oscillations which are transmitted to ON-dsRGCs, ultimately driving nystagmus. **D**, Pharmacological intervention in CSNB (*Nyx^nob^*) with DA: Application of DA increases retinal DA levels shifting the retina to its photopic wiring. This exogenous DA reduces the sensitivity of the A_II_ AC initial segments and likely decreases their coupling. As a result, A_II_ ACs are stabilized, their spontaneous oscillations cease.

CSNB mice suffer from low levels of DA in their retina due to defects in their rod ON-pathway, which is known to regulate DA release (26–29). Based on our results, we propose that this loss of DA input to the A_II_ ACs overly sensitizes their voltage-gated ion-channels, making these neurons prone to spontaneous oscillations (Figure 5C). Since these oscillations eventually cause nystagmus, our findings provide a cellular and molecular mechanism linking the retinal defect in CSNB to the manifestation of nystagmus.

DA is known to modulate the gap-junctions between A_II_ ACs and between A_II_ ACs and ON-BCs (21). Could this modulation account for our results? Since DA closes the gap-junctions, one would expect that the cell becomes more compact. The consequence of strong electrical coupling is that it increases the time constants of the measured currents. So, closing the gap-junctions would lead to a decrease instead of the increase in time constants we found. Therefore, our results cannot be accounted for by changes in gap-junction conductance alone. Furthermore, DA did not affect the resting membrane potential, which excludes the possibility that sustained hyperpolarization inhibited the oscillatory behavior of the A_II_ ACs. This leaves the modulation of the voltage-gated ion channels as the likely source of the change in oscillatory behavior. Indeed, DA can, depending on the concentration, increase (low concentration) or reduce (high concentration) the max conductance of potassium channels (31). We found a reduction in the peak current and a slowing down of I_K-A_. With regard to modulation of the I_Na_ channel, we found that the maximum conductance can be reduced by D1 DA receptor activation, which is similar to what Veruki and colleagues (2025) found in A_II_ ACs in rat retina and what others observed in hippocampal pyramidal neurons (32). Likewise, we found that the spikelets are attenuated in amplitude and pace by DA, which is in line with findings by Hayashida and colleagues (2009) in RGCs of rats (33).

Our model simulations indicate that the changes we observe in the voltage-gated channels in the A_II_ ACs are sufficient to account for the change in oscillatory behavior of the A_II_ ACs and thus for the modulation of the RGC oscillations and nystagmus. Still, DA can in principle also affect other processes and cells in the retina (21), and we can therefore not exclude the possibility that the light-response properties of the RGCs are also affected by dopaminergic modulation of one or more of the other retinal neurons. Our results support the hypothesis that dopamine acting via a D1 receptor on A_II_ ACs changes the oscillatory behavior of these neurons, while the signal strength seems to be affected by activation of both D1 and D2 receptors on other retinal neurons, including RGCs.

CSNB patients often suffer from poor visual performance (34). Our results suggest a possible cause for this. Suppressing the spontaneous oscillations markedly improved the SNR of RGC light responses in *Nyx^nob^* mice, which most likely has direct implications for visual performance. While developmental factors such as desegregation of retino-thalamic axons in the dorsolateral geniculate nucleus may affect the visual performance (35), our results suggest that low fidelity of information transfer at the retinal level, caused by the persistent background oscillations and low signal strength, forms a significant contributing factor. By reducing the oscillations (noise) and enhancing the amplitude of the light responses (signal), DA application not only blocks the oscillatory eye movements but also increases the SNR of the retinal output, offering a dual benefit for both stabilizing gaze and improving visual function.

Could our findings also offer insight into the cause of myopia, a common symptom in CSNB patients? Goethals and colleagues (36) have proposed that local contrast detector neurons in the retina may signal signs of defocus and thereby regulate growth of the eyes. Other mechanisms to detect defocus, such as longitudinal chromatic aberration and monochromatic aberration, have been suggested as well (37). Yet, independent of the mechanism that detects defocus, activity in the neurons performing this task has been consistently proposed to stimulate eye growth. Since flickering light has been identified as a trigger for axial elongation of the eye (38–40), it is therefore tempting to consider the possibility that in CSNB the oscillatory activity in the defocus-detecting neurons may promote eye growth, leading to myopia.

The finding that oscillating A_II_ ACs appear to be the main driver of the nystagmus associated with CSNB raises the question as to whether A_II_ AC instability could be a common underlying mechanism for nystagmus in a broader range of syndromes. A_II_ ACs are neurons that relay rod signals to the RGCs. A delicate balance between the activity of the voltage-gated ion channels on the initial segment, the DA release, and the membrane potential of the A_II_ ACs is needed to keep the A_II_ ACs stable. In many diseases this balance may be disturbed. A_II_ ACs are highly interconnected (Figures 5 and S1). Their main input is a glutamatergic input from ON rod-BCs, but they also receive inhibitory and neuromodulatory inputs, and they are also electrically coupled with other A_II_ ACs and cone ON-BCs. We hypothesize that any substantial depolarization of the A_II_ AC resting membrane potential due to a change in their synaptic input or disturbance in the dopaminergic input of the A_II_ ACs will lead to oscillations of A_II_ ACs and consequently RGCs and thereby to nystagmus. Several syndromes present with nystagmus and involve disruptions that could impact the delicate A_II_ AC balance. In albinism, the retinal dopaminergic system may be disturbed (41), which may have reduced the dopaminergic inputs to the A_II_ ACs and upregulated the voltage-gated ion channels, which could lead to oscillations and nystagmus. Similarly, nystagmus in Parkinson’s disease (42) may be directly due to reduced dopaminergic inputs to the A_II_ ACs. In early-onset severe retinal dystrophies, nystagmus could arise from disturbed glutamatergic inputs to the ON-BCs leading to depolarization of the A_II_ ACs (20). Moreover, observations in a mouse model of multiple sclerosis, indicating reduced photoreceptor synaptic output (43–45), also point towards aberrations in A_II_ AC depolarization, oscillations, and nystagmus.

The results presented in this paper could potentially lead to the first noninvasive, pharmacological treatment targeting infantile nystagmus associated with CSNB. Given that activation of D1 receptors blocks the A_II_ ACs and RGC oscillations and increases the RGC signal fidelity, application of D1 agonists or a combination of D1 and D2 agonists via eye-drops might form a potential treatment of nystagmus. Even though such treatment will not cure CSNB, it may dramatically reduce the burden of some of the symptoms, such as nystagmus, low visual performance, and possibly even myopia. The absence of rod vision will remain, but vision at the cone threshold may improve because of the increased SNR of the RGC signals. While validation in other infantile nystagmus models and long-term efficacy studies is necessary, this work provides significant mechanistic understanding into infantile nystagmus pathophysiology and presents topical dopaminergic activation—specifically D1 agonists—as a promising and testable therapeutic strategy, warranting further preclinical investigation.

## Materials and methods

### Animals

All animal experiments were carried out under the responsibility of the ethical committee of the Royal Netherlands Academy of Arts and Sciences (KNAW) acting in accordance with the European Communities Council Directive of 22 September 2010 (2010/63/EU). The experiments were performed under the license numbers AVD-801002016517 and AVD-80100202115698, issued by the Central Committee for Animal Experiments of the Netherlands.

*Nyx^nob^* mice were obtained from the McCall lab (University of Louisville, Louisville, USA). *Nyx^nob^* and WT mice were in a C57BL/6JRj background. Since the Nyx mutation is x-linked, only male mice in the age range of 5-71 weeks were used for the experiments. Room lights were timed on a 12/12 h light-dark schedule, and experiments took place during daytime. Mice had *ad libitum* access to food and water.

### Multielectrode RGC recordings

#### Retinal dissection

After 1 hour of dark adaptation, mice were sedated using a mixture of CO_2_/O_2_ and ultimately euthanized by cervical dislocation. All procedures were carried out under dim red light. The eyes were extracted from the eye socket and placed in room-temperature Ames’ medium (Sigma-Aldrich, St Louis, MO). Next, the cornea and lens were removed by making an insertion around the *ora serrata* using fine spring scissors. As much vitreous humor as possible, as well as the sclera was removed using fine forceps. Four small insertions were made, and the retina was flat-mounted on a filter paper annulus (3 mm inner radius; 0.8 µm hydrophilic MCE MF-MilliporeTM membrane filter, Merck Millipore Ltd., Tullagreen, Ireland). The retina was then mounted photoreceptor cell side up on a perforated 60-electrode MEA chip (60pMEA200/30iR-Ti, Multichannel systems, Reutlingen, Germany) in a recording chamber mounted on a Nikon Optiphot-2 upright microscope and viewed under IR with an Olympus 2x objective and video camera (Abus TVCC 20530). During the experiment the retina was continuously superfused with Ames’ medium gassed with a mixture of O_2_ and CO_2_ at a pH of 7.4 and a temperature of 35-37°C at a flow rate of 1.5 ml/min. Before the recording started, an acclimatization period in the dark for a duration of 15 minutes was given to ensure stable recordings.

#### Data acquisition

The extracellular RGC activity (n = 536 units, N = 10 retinas/mice) was acquired using MC rack (Multichannel systems, Reutlingen, Germany) at a sampling frequency of 25 kHz. The data was then zero-phase bandpass filtered (250-6250 Hz) with a fourth-order Butterworth filter in Matlab (Mathworks, Natick, MA, USA). Subsequently, the spiking activity was sorted manually into single-unit activity using the Plexon offline sorter (Plexon, Dallas, TX, USA) based on the first two principal components versus time. For the extraction of spikes from the background, a spike detection amplitude threshold of > 4σ_n_ was used as criteria. Here, σ_n_ is defined as an estimation of the standard deviation of background noise and *x* is the bandpass-filtered signal (Quiroga *et al.*, 2004; Holzel *et al.*, 2023).

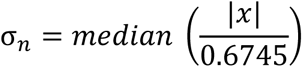

#### Optical stimulator

Full-field white light flashes were generated using Psychophysics Toolbox Version 3 (Brainard, 1997) and presented to the photoreceptor side of the retina with a custom-modified DLP projector (custom modification(46); Light Crafter 4500, Wintech, Carlsbad, CA, USA). Only white light stimuli were used. White light stimuli consisted of equal quantal output of the red (625 nm), green (530 nm),blue (455 nm), and UV (385 nm) LEDs. The maximal light intensity was 8.60 × 10^16^ quanta m^−2^ s^−1^.

#### Data analysis

To quantify the oscillatory behavior of RGC’s activity, the firing activity in the dark was recorded for a period of at least 15 minutes, cut into 5s non-overlapping epochs. The setup produces a series of spike times; this data was converted into a time series with 1 ms resolution. Each epoch was detrended and baseline subtracted, after which the autocorrelation and Welch’s PSD were calculated for each cell per trace and then averaged per cell.

For the signal-to-noise analysis, PSTHs were made from each cell’s activity in 10 ms bins. The signal-to-noise ratio was defined as the response divided by the STD. The response was defined as the absolute deviation from the mean after light offset, as we are only analyzing the OFF response due to the absence of an ON response in the *Nyx^nob^* mice. The mean was taken in the dark period measured before the stimulus. The noise was determined from a long dark measurement taken before the stimulus protocol, after first synchronizing the oscillatory activity in each trace to the first burst, after which PSTHs were made and the STD was calculated. We synchronized the bursting activity to have a more fair measure of the variance in the signal since summing unsynchronized oscillatory activity will be turned into a stable but higher constant signal (as seen in Figure 4D). The synchronized activity seen in the PSTH still dampens out over time due to small variation in the frequency. As a control, we did run the analysis on the unsynchronized data as well and still found an increase in the SNR.

Time constants were determined by fitting a double exponential curve through the voltage clamp data.

### Ideal observer

We investigated the effect of dopamine on the signal quality with an ideal observer type of analysis. For this analysis we stimulated the retina with 3 groups of 2 stimuli that differed in intensity by 5%. The intensity of the groups differed by 0.33 log units. These stimuli were repeated 100 times. The data was split into 2 datasets consisting of the even and the uneven traces from which PSTHs were made. The even traces were used as data, and the uneven traces to generate masks of the responses per cell. The masks were created by fitting straight lines through segments of the dataset. We chose the segments based on the steady and dynamic parts of the response. The steady parts, 0 to 550 ms, 1500 to 1600 ms, and 1600 to 2000 ms, were fitted with single straight lines. The dynamic parts, 550 to 650 and 1050 to 1500, were fitted with a line between every 3 data points. Finally, we took the sum of the squared errors between the response and the mask to determine which mask fits the response best and used that as a quantity representing the ability to discriminate between the 2 stimuli.

### Statistical analysis

Statistical analyses were performed using GraphPad Prism 10.0.3 (GraphPad Software, Massachusetts, USA) or Matlab (MathWorks Inc., Natick, MA). All mean values are presented ± SEM. Statistical significance was tested using a Wilcoxon matched-paired signed rank test for paired data and a Mann-Whitney test for unpaired data. Normality was tested using the Shapiro-Wilk test.

### Whole cell A_II_ AC recordings

For recordings of A_II_ ACs (n = 16 cells, N = 12 mice), vertical slices of mouse retina (~200 µm) were prepared using a custom-made tissue slicer. Animals were dark-adapted for 1 hour. Anesthetization and all subsequent procedures were performed under infrared illumination to preserve the adaptation state. During recordings, cells were visualized and targeted under infrared illumination. Slices were transferred to the recording chamber and perfused at a rate of ~2 ml/min with bicarbonate-buffered Ames medium (Sigma-Aldrich) continuously bubbled with 95% O2, 5% CO2 at pH of 7.4. Slices were visualized with a Nikon Eclipse E2000FN microscope fitted with a 60x/1.0 water immersion objective with an infrared differential interference contrast, a video camera (Philips, The Netherlands) and a two-photon scan head. A_II_ ACs were identified by their location, morphology, and characteristic current responses. Recordings were performed at 35-37 °C. Series resistance, which was always below 30 MΩ, was not compensated, and the liquid junction potential was not corrected.

Patch electrodes were fabricated from thick-walled borosilicate glass to have a resistance of 5-7 MΩ. The intracellular solution contained (in mM): K-methanesulfonate, 135; KCl, 6; Na2-ATP, 2; Na-GTP, 1; EGTA, 1; MgCl2, 2; Na-HEPES, 5; and adjusted to pH 7.35 with KOH. Lucifer yellow (1 mg/ml) was added to these solutions for morphological identification of recorded cells using a two-photon microscope. Whole-cell recordings were made with a Multiclamp 700B amplifier using a Digidata 1322A digitizer (MDS Analytical Technologies, Union City, CA) and Signal 6.04 software (Cambridge Electronic Design Ltd, Cambridge, UK) to generate command outputs, trigger the light stimulus, and acquire and analyze analog whole-cell voltages. These data were sampled at 10 kHz and filtered at 2.4 kHz with a four-pole Bessel low-pass filter. Matlab (MathWorks Inc., Natick, MA) and Igor Pro (WaveMetrics, Portland, OR, USA) were used to analyze the data.

### Computational model

Figure 3A illustrates the simulated network composed of an ON-BC, A_II_ AC, and RGC. In this network, A_II_ AC form electrical connections via gap junctions with cone-driven ON-BC (ON-CBC). The A_II_ AC receives excitatory glutamatergic input from ON-RBCs(14) and inhibitory input from OFF-ACs that are activated by OFF-CBCs(14). This inhibitory connection represents a form of crossover inhibition. When light is presented, ON BCs depolarize, which consequently depolarizes the A_II_ ACs. At the same time, light stimulation causes OFF-CBCs and thus the OFF-ACs they drive to hyperpolarize, reducing their inhibitory influence on A_II_ ACs and further enhancing A_II_ AC depolarization. ON-CBCs transmit signals to ON-RGCs, whereas OFF-BCs project to OFF-RGCs. In the model there is only one ON-CBC per A_II_ AC although literature shows that one A_II_ AC is contacted by 10 ON-CBCs(47). This physiological connectivity was implemented by scaling the coupling conductance. Next, we present the equations describing the various cell types.

#### A_II_ AC model

The A_II_ AC was modeled using a three-compartment framework adapted from Choi et al., 2014. This model includes a soma, an initial segment, and an interconnecting cable. The dendritic arbor is represented as part of the somatic compartment. All ionic currents in the model are governed by Hodgkin-Huxley-type equations.

The sodium current (*I_Na_*) is given by

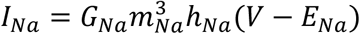

where the gating variables are

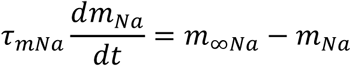

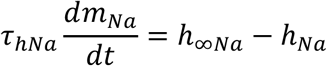

with

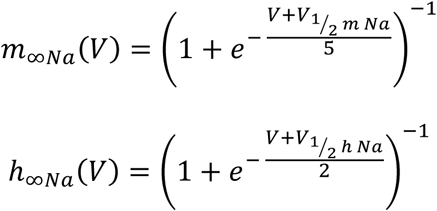

The M-type potassium current (*I_M_*) is given by

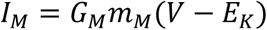

where the gating variable are

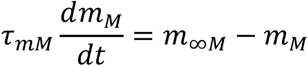

with

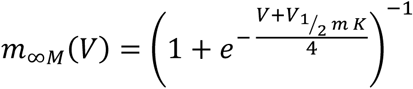

The A-type potassium current (*I_A_*) is given by

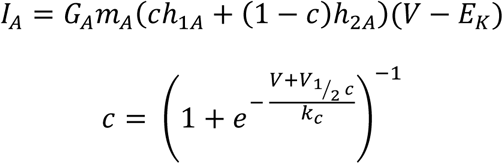

where the gating variables are

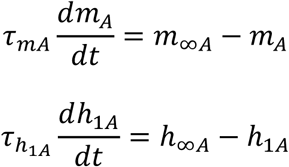

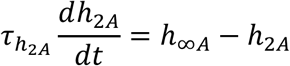

with

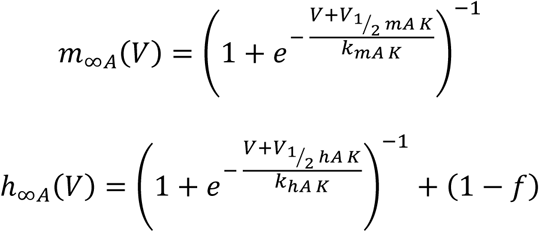

Next to these ionic currents there is a leak current

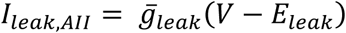

The ON-CBCs are connected to the A_II_ ACs through gap junctions. For computational efficiency, each A_II_ AC was coupled with a single ON-CBC in the simulation. To represent the biological condition in which one A_II_ AC is coupled to approximately 10 ON-CBCs, the gap junction conductance on the A_II_ AC side was set to be 10 times greater than that experienced by the ON-CBC.

The gap-junction currents are given by

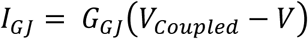

The membrane potentials of the three compartments were calculated as

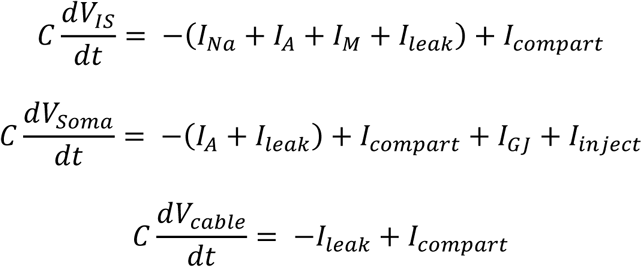

The model parameters are summarized in Table 1.

#### ON-CBC model

For the ON-CBC, we used the model from Usui *et al.* (1996). This model consists of a delayed rectifying potassium current (*I_Kv_*), a transient potassium current (*I_A_*), a hyperpolarization-activated current (*I_h_*), a calcium current (*I_Ca_*), and a calcium-dependent potassium current (*I_KCa_*). A two-compartment model was used for the internal calcium concentration of the ON-CBC, which is needed to calculate I_KCa_. One compartment was close to the membrane and consisted of a fast and a slow calcium buffer and two exchange currents. The other compartment was within the cell and consisted of only the two calcium buffers. Calcium will flow from one compartment into the other by diffusion.

All ionic currents are described with Hodgkin-Huxley-type equations, except for I_h_ which is described with a Markov chain-type model.

The delayed rectifying potassium current (*I_Kv_*) current is described by

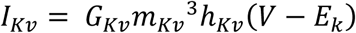

where the gating variables are calculated as

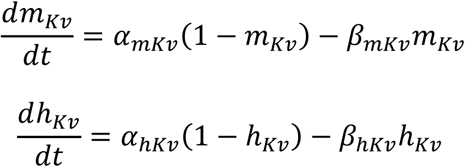

with

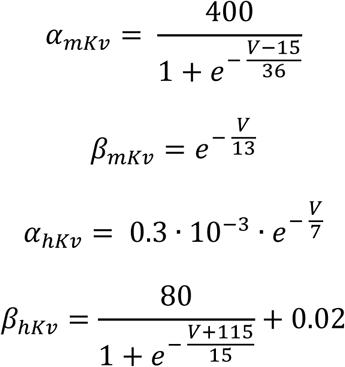

The transient potassium current (I_A_) is modeled with Hodgkin-Huxley type equations, and is defined as

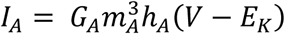

with the gating variables

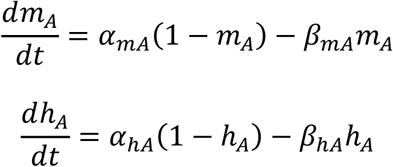

where

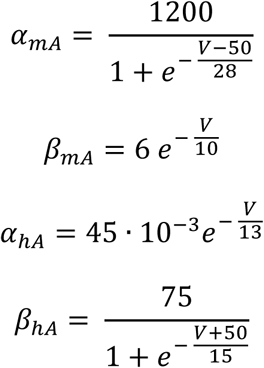

The hyperpolarization activated current (*I_h_*) is described with a Marko chain like model which described the transition between open and closed states. The current is given by

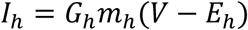

where the total open probability is the sum of the open states

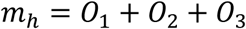

The open and closed states as calculated as

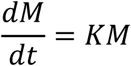

where

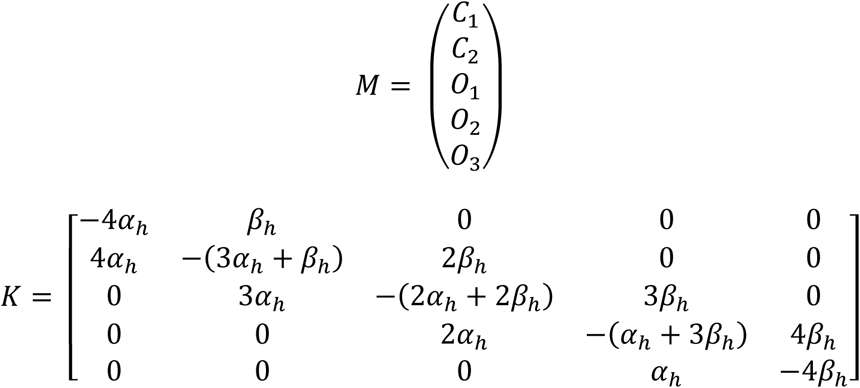

With the transition rate constants defined as

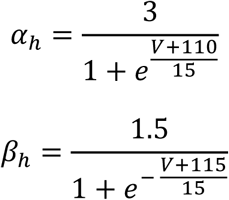

The calcium current (*I_Ca_*) is given by

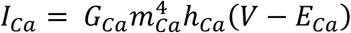

with a reversal potential depended on the calcium concentration

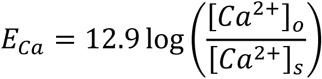

with the gating variables

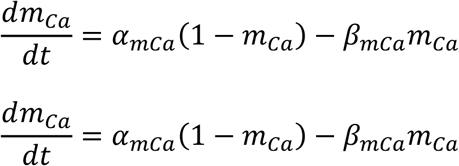

where

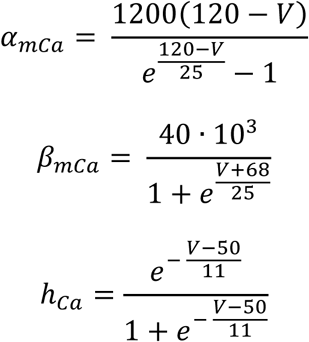

The calcium dependent potassium current (*I_KCa_*) is given by

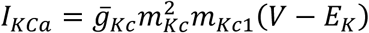

where the gating variables are described as

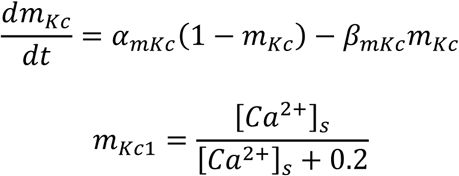

with

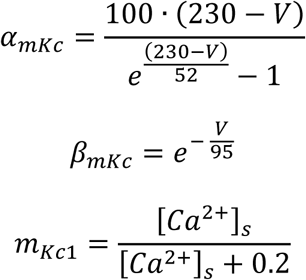

The change in calcium concentration close to the membrane ([Ca^+2^]_s_) is described as

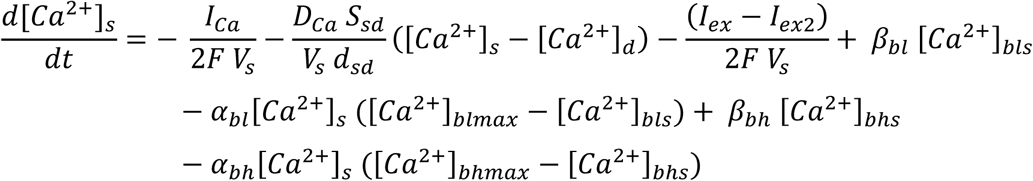

and the calcium concentration inside the cell ([Ca^2+^]_d_) as

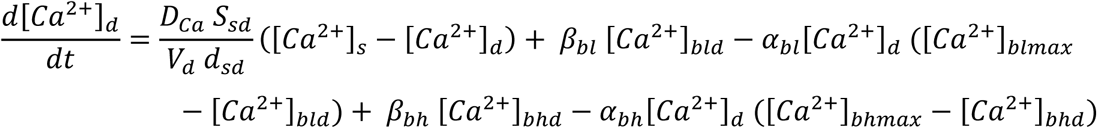

where

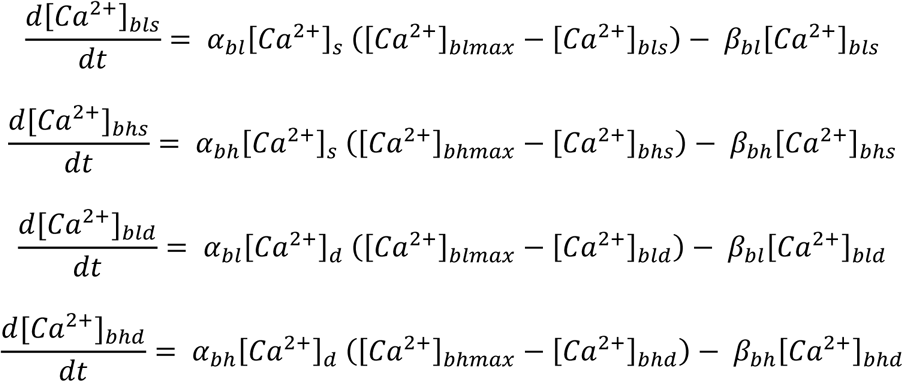

with the exchange currents defined as

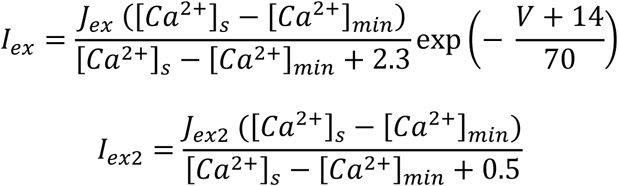

Finally, we added a leak current

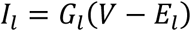

The gap junction between the ON-CBC and the A_II_ AC is given by.

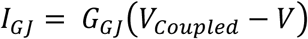

We only simulate one ON-CB per A_II_ cell. The membrane potential of the BC is given by

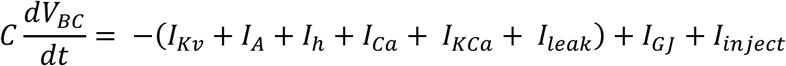

The model parameters are summarized in Tables 2 and 3.

**Table 2.** ON-CBC and RGC model parameters.

| ON-CBC |  |
| --- | --- |
| C | 10 pF |
| ***Delayed rectifying potassium current*** |  |
| $G_{Kv}$ | 2 nS |
| $E_{Kv}$ | -58 mV |
| ***The transient potassium current*** |  |
| $G_A$ | 35 nS |
| $E_K$ | -58 mV |
| ***Hyperpolarization activated current*** |  |
| $G_h$ | 5.975 nS |
| $E_h$ | -17.7 mV |
| ***Calcium current*** |  |
| $G_{Ca}$ | 1.1 |
| $[Ca^{2+}]_o$ | 2000 μM |
| ***Calcium dependent potassium current*** |  |
| $G_{Kc}$ | 8.5 nS |
| $E_K$ | -58 mV |
| ***Leak current*** |  |
| $G_l$ | 0.5 nS |
| $E_l$ | -50 mV |
| ***Gap-junction*** |  |
| $G_{GJ-A2-BC}$ | 20 nS |
| ON-RGC |  |
| C | $1.26 \times 10^{-10}$ S |
| $E_k$ | -77 mV |
| $E_{na}$ | 50 mV |
| $E_{leak}$ | -100 mV |
| $g_{leak}$ | $3.1 \times 10^{-12}$ S |
| $G_{syn}$ | $10 \times 10^{-12}$ S |

**Table 3.**
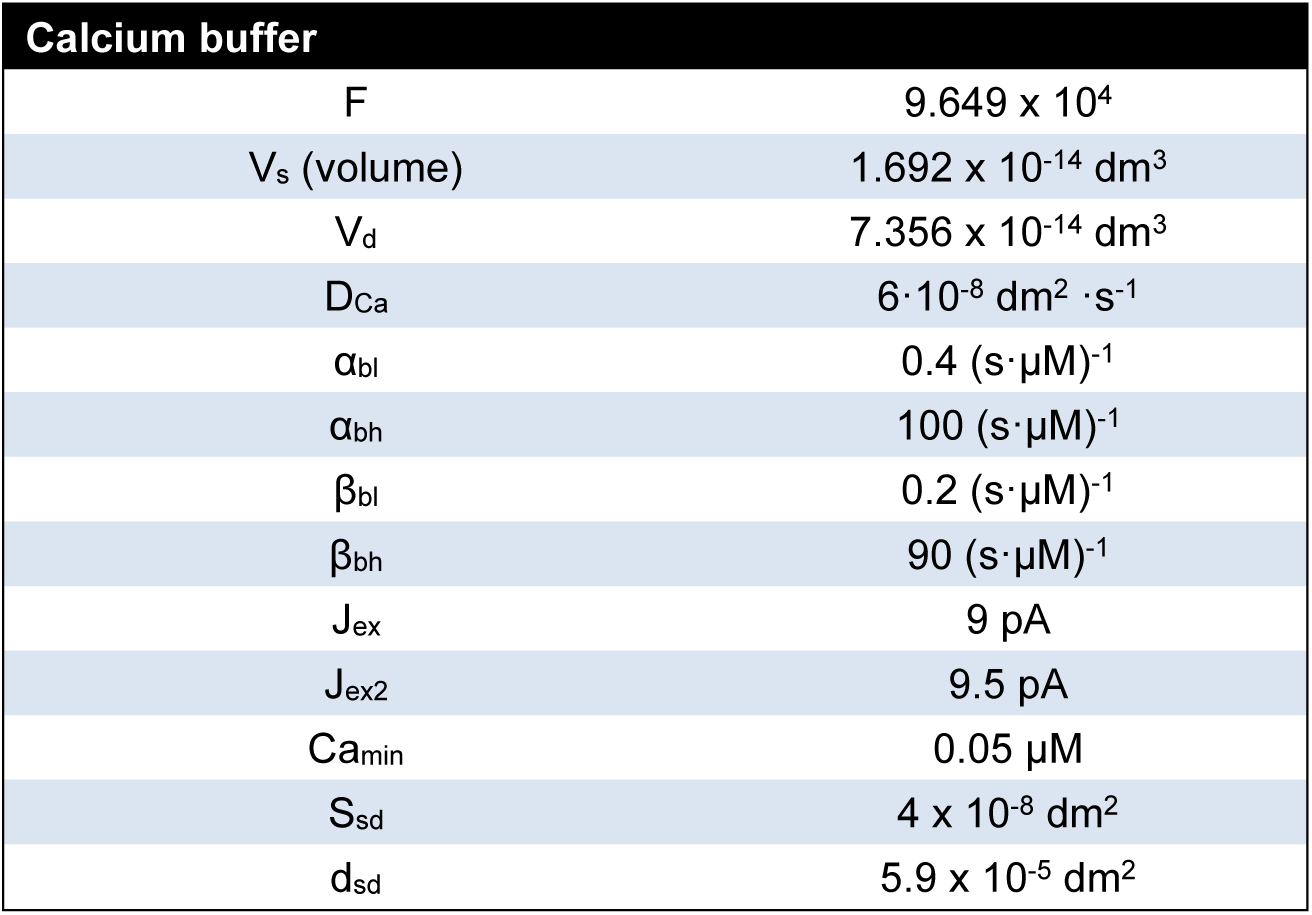
ON-CBC calcium buffer parameters.

#### RGC model

The RGC model is a simple spike-generating mechanism consisting of three ion channels, a sodium current, an M-Type potassium current, an A-Type potassium channel, and a leak current. In addition to these ion channels, there is synaptic current, which is modeled as a current that is dependent on the calcium current of the bipolar cell. The sodium channel, M-Type potassium channel, and A-Type potassium channels are described with the same equation used for the A_II_ ACs. The synaptic current is given by

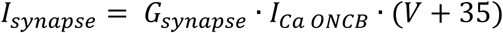

The membrane potential of the RGC is given by

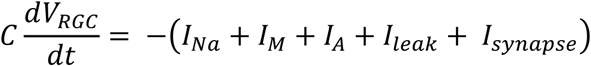

## Acknowledgement

We thank Cynthia Geelen for her technical support. This work was supported by grants from Uitzicht (2023-07; MK), Van Hessen-Israels fund (MK), Dutch Organization for Medical Sciences (ZonMW 91120067; MK and CIDZ), INTENSE LSH-NWO (TTW/00798883, CIDZ) and NWO-Gravitation (DBI2 024.005.022; CIDZ).

## Author contributions

BSS, Experiments, analysis, writing, conceptualization.

WK, Model simulation, analysis, writing, conceptualization.

BW, Conceptualization.

CIDZ, Funding, writing, secondary conceptualization.

MK, Funding, writing, conceptualization.

## Supplementary Information

**Figure S1.**
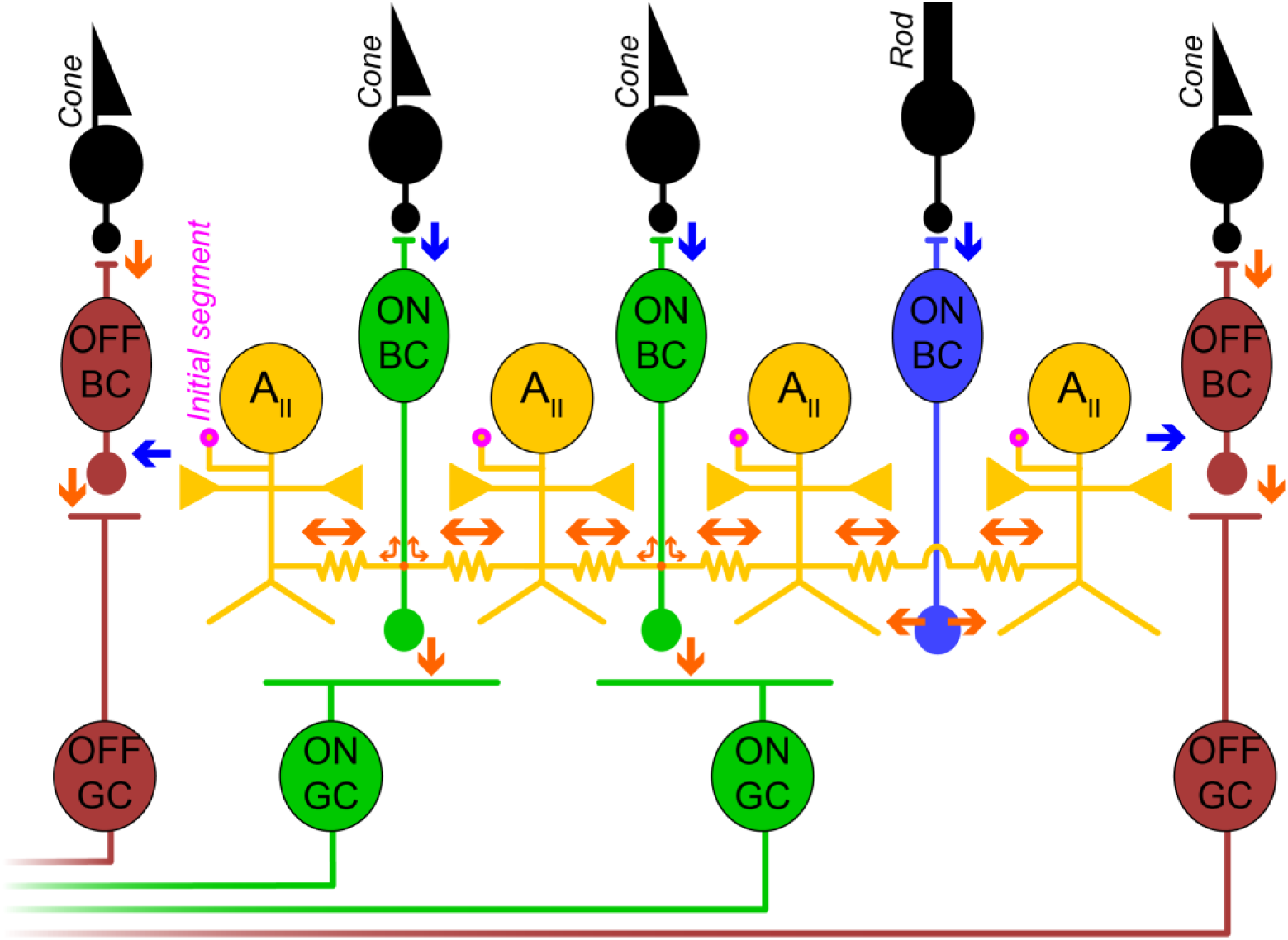
Schematic of retinal microcircuit highlighting A_II_ AC interconnectivity. The diagram illustrates a retinal microcircuit comprising photoreceptors (cones and a rod), bipolar cells (OFF cone BCs in maroon, ON cone BCs in green, and a rod ON BC in blue), A_II_ ACs (yellow), and ganglion cells (OFF GCs in maroon, ON GCs in green). The extensive interconnectivity of the A_II_ ACs includes: (i) Direct electrical coupling between A_II_ ACs via gap junctions (indicated by double-headed orange arrows), forming a network of these cells. (ii) Chemical synaptic input received from rod ON BCs (blue arrow from the rod ON BC to an A_II_ AC). (iii) Electrical coupling with ON cone-BCs via gap junctions, allowing signals from the rod pathway to flow into the ON cone pathway. (iv) Chemical synaptic output to OFF cone BCs (blue arrows from A_II_ ACs to OFF cone BCs), which relays rod-driven signals to the OFF pathway. The “Initial segment” of an A_II_ AC, a site critical for its electrophysiological properties, is labeled in magenta. Blue arrows represent sign-inverting pathways, and red arrows represent sign-conserving pathways.

**Figure S2.**
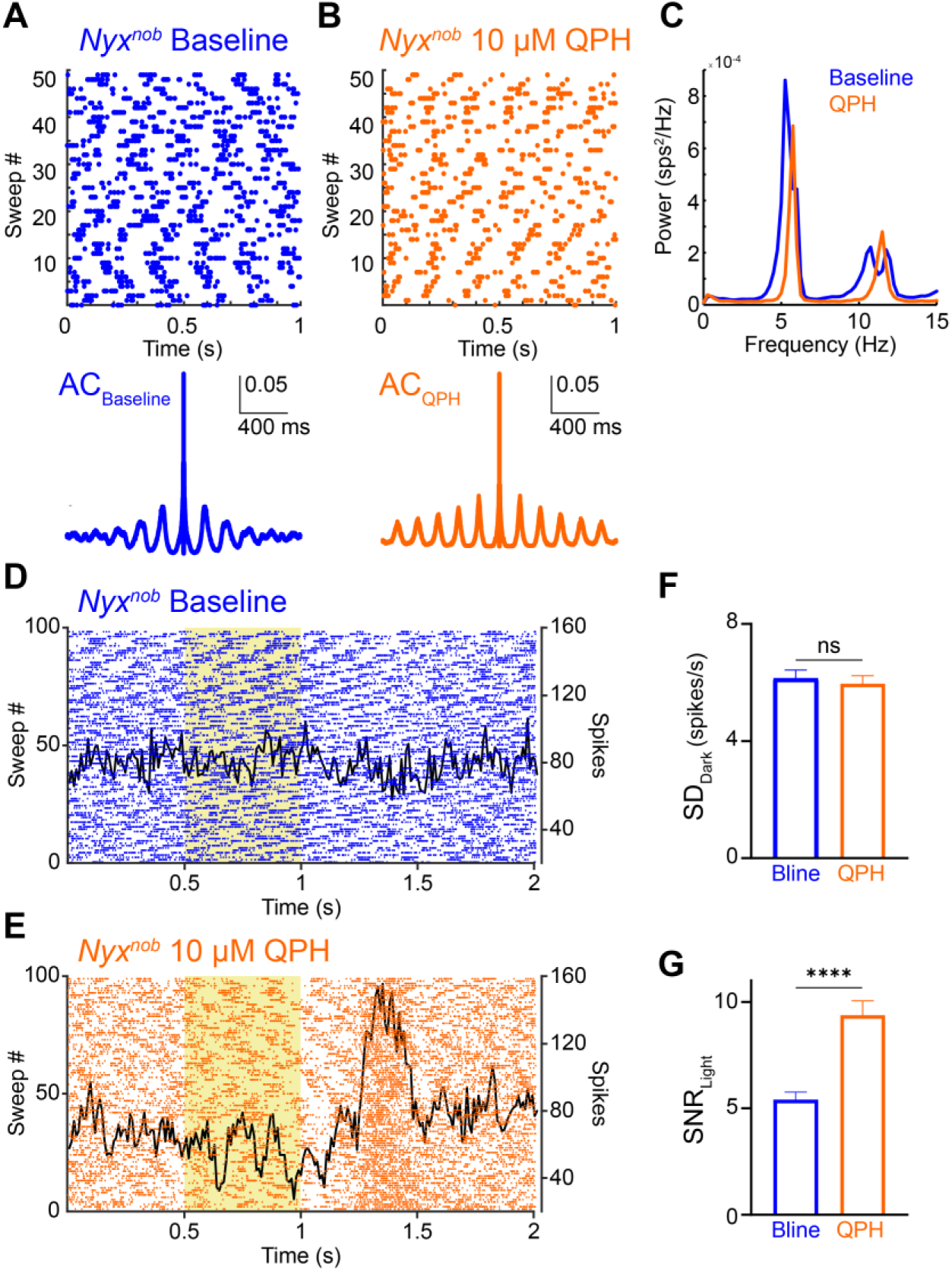
Activation of D2-receptor failed to eliminate oscillations but enhanced the signal-to-noise ratio. **A-B**, Spontaneous retinal ganglion cell (RGC) activity in the dark in an isolated light-adapted retina. Top: raster plot of a single RGC. Bottom: autocorrelation of the same RGC. **A**, Baseline *Nyx^nob^* mouse retina showing robust oscillations. **B,** *Nyx^nob^* retina with 10 µM quinpirole hydrochloride (QPH) application. **C**, Mean power spectral density of RGC activity from a representative retina under baseline conditions (blue) and with QPH (red). **D**, Example of a single RGC’s response to light stimulation in a baseline *Nyx^nob^* retina. Raster plot (blue). Overlaid corresponding peri-stimulus time histogram (PSTH, black). **E**, Same single RGC as in (**D**) but after application of 10 µM QPH (orange), illustrating increased peak light response. **F**, Bar graphs quantifying the mean standard deviation (SD) of baseline RGC firing rates (noise) in the dark period preceding light stimulation under baseline (blue) and 10 µM QPH (orange) conditions. SD: baseline (blue; mean 6.14, n = 122) vs QPH (orange: 5.96, n = 122; *P* = 0.23, Wilcoxon test) **G**, Bar graphs quantifying the mean signal-to-noise ratio (SNR) of RGC light responses for baseline (blue) and 10 µM QPH (orange) conditions. SNR: baseline (blue; mean 5.39, n = 122) vs QPH (orange: 9.38, n = 122; *P* < 0.0001, Wilcoxon test).

**Figure S3.**
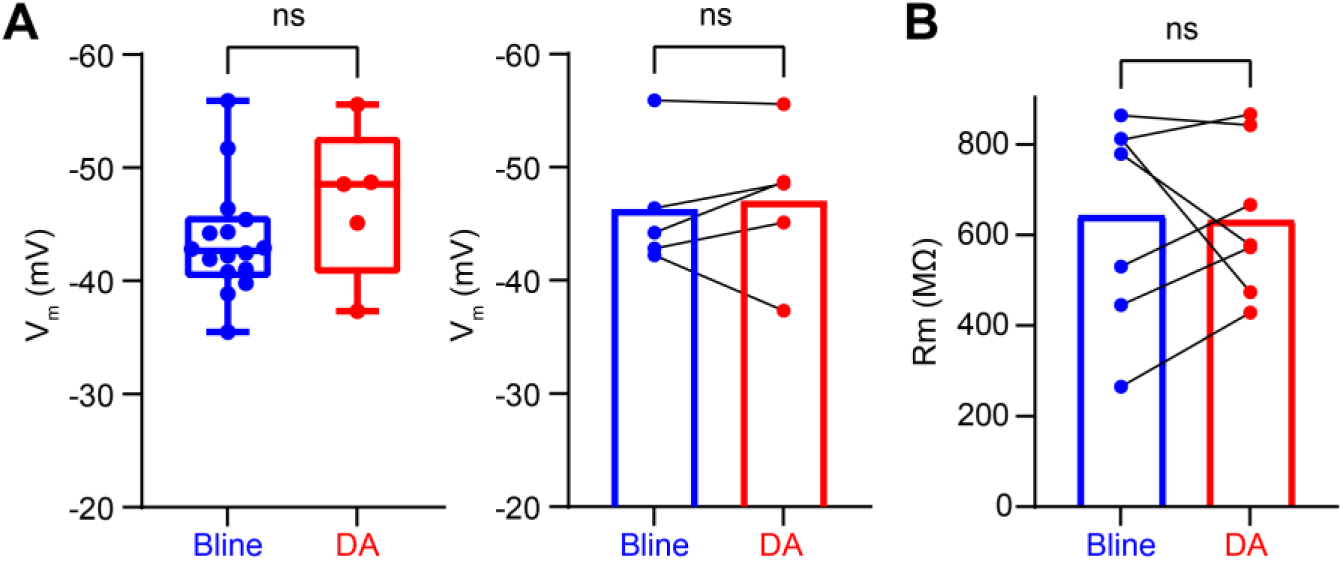
Dopamine does not alter passive membrane properties. **A**, Left: Box plots show the distribution of V_m_ (mV) under baseline (Bline, blue) and 10 µM DA (red) conditions. V_m_: baseline (blue; mean −43.5 mV, n = 16) vs DA (red; mean −47.1 mV, n = 5; *P* = 0.21, Mann-Whitney test). Right: Bar graphs with paired circles show data from individual cells. V_m_: baseline (blue; mean −46.3 mV, n = 5) vs DA (red; mean −47.1 mV, n = 5; *P* = 0.81, Wilcoxon test) **B**, Bar graphs show the mean membrane resistance (R_m_, MΩ) under baseline (Bline, blue) and 10 µM DA (red) conditions. Paired circles show data from individual cells. R_m_: baseline (blue; mean 644.1 MΩ, n = 7) vs DA (red; mean 632.8 MΩ, n = 7; *P* > 0.999, Wilcoxon test).

## References

1. J. I. Simpson, D. R. W. Wylie, C. I. D. Zeeuw, More on Climbing Fiber Signals and Their Consequence(S). Behavioral and Brain Sciences 19, 496–498 (1996).

2. D. S. Zee, R. J. Leigh, Disorders of Eye Movements. Neurologic Clinics 1, 909–928 (1983).

3. L. F. Dell’Osso, Grating visual acuity in infantile nystagmus in the absence of image motion. Invest Ophthalmol Vis Sci 55, 4952–4954 (2014).

4. M. J. Dunn, et al., Grating visual acuity in infantile nystagmus in the absence of image motion. Invest Ophthalmol Vis Sci 55, 2682–2686 (2014).

5. J. Felius, et al., Quantifying nystagmus in infants and young children: relation between foveation and visual acuity deficit. Invest Ophthalmol Vis Sci 52, 8724–8731 (2011).

6. J. Felius, Z. A. Muhanna, Visual Deprivation and Foveation Characteristics Both Underlie Visual Acuity Deficits in Idiopathic Infantile Nystagmus. Invest. Ophthalmol. Vis. Sci. 54, 3520–3525 (2013).

7. H. J. Simonsz, R. J. Florijn, H. M. van Minderhout, A. A. B. Bergen, M. Kamermans, Nightblindness-Associated Transient Tonic Downgaze (NATTD) in Infant Boys with Chin-Up Head Posture. Strabismus 17, 158–164 (2009).

8. J. Klooster, et al., Ultrastructural Localization of GPR179 and the Impact of Mutant Forms on Retinal Function in CSNB1 Patients and a Mouse Model. Invest. Ophthalmol. Vis. Sci. 54, 6973–6981 (2013).

9. N. T. Bech-Hansen, et al., Mutations in NYX, encoding the leucine-rich proteoglycan nyctalopin, cause X-linked complete congenital stationary night blindness. Nat Genet 26, 319–323 (2000).

10. R. G. Gregg, et al., Nyctalopin expression in retinal bipolar cells restores visual function in a mouse model of complete X-linked congenital stationary night blindness. J Neurophysiol 98, 3023–3033 (2007).

11. M.-B. Hölzel, et al., A common cause for nystagmus in different congenital stationary night blindness mouse models. J Physiol (2023). 10.1113/JP284965.

12. M. Kamermans, et al., A retinal origin of nystagmus-a perspective. Front Ophthalmol (Lausanne) 3, 1186280 (2023).

13. B. H. J. Winkelman, et al., Nystagmus in patients with congenital stationary night blindness (CSNB) originates from synchronously firing retinal ganglion cells. PLOS Biology 17, e3000174 (2019).

14. R. E. Marc, J. R. Anderson, B. W. Jones, C. L. Sigulinsky, J. S. Lauritzen, The AII amacrine cell connectome: a dense network hub. Front Neural Circuits 8, 104 (2014).

15. R. G. Smith, N. Vardi, Simulation of the AII amacrine cell of mammalian retina: functional consequences of electrical coupling and regenerative membrane properties. Vis Neurosci 12, 851–860 (1995).

16. B. Völgyi, M. R. Deans, D. L. Paul, S. A. Bloomfield, Convergence and Segregation of the Multiple Rod Pathways in Mammalian Retina. J. Neurosci. 24, 11182–11192 (2004).

17. M. L. Veruki, J. H. Liu, J. B. Singh, M. S. Luppi, E. Hartveit, Activation of Dopamine D1 Receptors at the Axon Initial Segment-Like Process of Retinal AII Amacrine Cells Modulates Action Potential Firing. J Neurosci 45, e0736252025 (2025).

18. M. S. Cembrowski, et al., The mechanisms of repetitive spike generation in an axonless retinal interneuron. Cell Rep 1, 155–166 (2012).

19. D. J. Margolis, A. J. Gartland, J. H. Singer, P. B. Detwiler, Network oscillations drive correlated spiking of ON and OFF ganglion cells in the rd1 mouse model of retinal degeneration. PLoS One 9, e86253 (2014).

20. H. Choi, et al., Intrinsic bursting of AII amacrine cells underlies oscillations in the rd1 mouse retina. Journal of Neurophysiology 112, 1491–1504 (2014).

21. S. Roy, G. D. Field, Dopaminergic modulation of retinal processing from starlight to sunlight. J Pharmacol Sci 140, 86–93 (2019).

22. M. Contini, E. Raviola, GABAergic synapses made by a retinal dopaminergic neuron. Proceedings of the National Academy of Sciences 100, 1358–1363 (2003).

23. J. R. Anderson, et al., Exploring the retinal connectome. Mol Vis 17, 355–379 (2011).

24. G. Debertin, et al., Tyrosine hydroxylase positive perisomatic rings are formed around various amacrine cell types in the mammalian retina. J Neurochem 134, 416–428 (2015).

25. P. S. Junior, C. M. Wakeham, H. von Gersdorff, Dopamine regulates the membrane potential and glycine release of AII amacrine cells via D1-like receptor modulation of gap junction coupling. [Preprint] (2024). Available at: https://www.biorxiv.org/content/10.1101/2024.12.11.625486v1 [Accessed 23 October 2025].

26. M. H. Aung, et al., ON than OFF pathway disruption leads to greater deficits in visual function and retinal dopamine signaling. Experimental Eye Research 220, 109091 (2022).

27. M. T. Pardue, et al., High Susceptibility to Experimental Myopia in a Mouse Model with a Retinal ON Pathway Defect. Investigative Ophthalmology & Visual Science 49, 706–712 (2008).

28. T. Munteanu, et al., Light-dependent pathways for dopaminergic amacrine cell development and function. Elife 7, e39866 (2018).

29. V. Pérez-Fernández, et al., Rod Photoreceptor Activation Alone Defines the Release of Dopamine in the Retina. Curr Biol 29, 763–774.e5 (2019).

30. M. Tian, T. Jarsky, G. J. Murphy, F. Rieke, J. H. Singer, Voltage-Gated Na Channels in AII Amacrine Cells Accelerate Scotopic Light Responses Mediated by the Rod Bipolar Cell Pathway. J Neurosci 30, 4650–4659 (2010).

31. S. F. Fan, S. Yazulla, Modulation of voltage-dependent K+ currents (IK(V)) in retinal bipolar cells by ascorbate is mediated by dopamine D1 receptors. Vis Neurosci 16, 923–931 (1999).

32. A. R. Cantrell, R. D. Smith, A. L. Goldin, T. Scheuer, W. A. Catterall, Dopaminergic modulation of sodium current in hippocampal neurons via cAMP-dependent phosphorylation of specific sites in the sodium channel alpha subunit. J Neurosci 17, 7330–7338 (1997).

33. Y. Hayashida, et al., Inhibition of Adult Rat Retinal Ganglion Cells by D1-Type Dopamine Receptor Activation. J Neurosci 29, 15001–15016 (2009).

34. M. M. C. Bijveld, et al., Genotype and phenotype of 101 dutch patients with congenital stationary night blindness. Ophthalmology 120, 2072–2081 (2013).

35. J. Demas, et al., Failure to maintain eye-specific segregation in nob, a mutant with abnormally patterned retinal activity. Neuron 50, 247–259 (2006).

36. S. Goethals, et al., Nonlinear spatial integration allows the retina to detect the sign of defocus in natural scenes. Sci. Adv. 11, eadq6320 (2025).

37. F. Schaeffel, C. Wildsoet, Can the retina alone detect the sign of defocus? Ophthalmic and Physiological Optics 33, 362–367 (2013).

38. Y. Di, et al., Effects of Chronic Exposure to 0.5 Hz and 5 Hz Flickering Illumination on the Eye Growth of Guinea Pigs. Current Eye Research (2013).

39. J. Tang, et al., Effects of 2 Hz flickering light on refractive state, fundus imaging and visual function of C57BL/6 mice. Exp Eye Res 246, 110014 (2024).

40. Y. Yu, H. Chen, J. Tuo, Y. Zhu, Effects of flickering light on refraction and changes in eye axial length of C57BL/6 mice. Ophthalmic Res 46, 80–87 (2011).

41. A. Sanchez-Bretano, et al., Human equivalent doses of L-DOPA rescues retinal morphology and visual function in a murine model of albinism. Sci Rep 13, 17173 (2023).

42. I. Jung, J.-S. Kim, Abnormal Eye Movements in Parkinsonism and Movement Disorders. J Mov Disord 12, 1–13 (2019).

43. E. Dembla, M. Dembla, S. Maxeiner, F. Schmitz, Synaptic ribbons foster active zone stability and illumination-dependent active zone enrichment of RIM2 and Cav1.4 in photoreceptor synapses. Sci Rep 10, 5957 (2020).

44. A. Kesharwani, K. Schwarz, E. Dembla, M. Dembla, F. Schmitz, Early Changes in Exo- and Endocytosis in the EAE Mouse Model of Multiple Sclerosis Correlate with Decreased Synaptic Ribbon Size and Reduced Ribbon-Associated Vesicle Pools in Rod Photoreceptor Synapses. Int J Mol Sci 22, 10789 (2021).

45. A. Mukherjee, et al., Disturbed Presynaptic Ca2+ Signaling in Photoreceptors in the EAE Mouse Model of Multiple Sclerosis. iScience 23, 101830 (2020).

46. M.-B. Hölzel, M. H. C. Howlett, M. Kamermans, Receptive Field Sizes of Nyxnob Mouse Retinal Ganglion Cells. Int J Mol Sci 23, 3202 (2022).

47. C. L. Sigulinsky, et al., Network Architecture of Gap Junctional Coupling among Parallel Processing Channels in the Mammalian Retina. J Neurosci 40, 4483–4511 (2020).

